# Epigenetic dysregulation of Th2 cytokine genes in MuSK myasthenia gravis and its modulation by immunosuppressive therapy

**DOI:** 10.64898/2026.08.05.742975

**Authors:** Cansu Elmas, Andrea Stoccoro, Martina Lari, Farzin Salehi, Veronica Iovino, Alba Cepele, Jennifer Huber, Francisca Faber, Marlene Wolfsgruber, Omar Keritam, Rosa Weng, Anja Steinmaurer, Theresa König, Melania Guida, Hakan Cetin, Fritz Zimprich, Romana Höftberger, Michelangelo Maestri Tassoni, Fabio Coppedè, Inga Koneczny

## Abstract

**Background and objectives:** Myasthenia gravis associated with antibodies against muscle-specific kinase (MuSK-MG) is a well-characterized IgG4-autoimmune disease, however, the mechanisms driving IgG4 predominance remain poorly understood. This study investigated whether promoter DNA methylation of cytokine genes involved in IgG4 class switching is associated with this immune response.

**Methods:** Peripheral blood mononuclear cells were isolated from MuSK-MG patients (n=36), acetylcholine receptor myasthenia gravis (AChR-MG) patients as disease controls (n=7), and sex-matched healthy controls (n=12). Promoter DNA methylation of *IL4*, *IL10*, and *IL13* was assessed by methylation-sensitive high-resolution melting and relative cytokine mRNA expression by qPCR. Associations with clinical variables, and antibody levels were subsequently evaluated.

**Results:** MuSK-MG patients showed lower median *IL13* promoter methylation compared with healthy controls (*p* = 0.004). Median *IL4* promoter methylation was also reduced in MuSK-MG compared with healthy controls (*p* < 0.001) and AChR-MG disease controls (*p* < 0.001), whereas no differences were observed for *IL10* promoter methylation. Relative mRNA expression of *IL4* (*p* = 0.0005), *IL10* (*p* = 0.0462), and *IL13* (*p* = 0.0002) was increased in MuSK-MG compared with AChR-MG. Compared with healthy controls, only *IL4* expression remained significantly increased (*p* < 0.0001). Promoter methylation was inversely correlated with relative mRNA expression for *IL4* (*p* < 0.0001), while *IL13* showed a similar but non-significant trend (*p* = 0.054), no association was observed for *IL10*. Multivariable analysis demonstrated that treatment at sampling was independently associated with lower *IL10* and *IL13* promoter methylation, whereas no associations were observed with age, sex, disease phase, or disease duration. Promoter methylation did not correlate with total serum IgG4 or anti-MuSK IgG4 levels.

**Discussion:** MuSK-MG is associated with selective hypomethylation of *IL4* and *IL13* promoters accompanied by increased cytokine gene expression, while *IL10* promoter methylation remains unchanged. The association between treatment and *IL10* and *IL13* promoter methylation suggests that immunosuppressive therapy may influence epigenetic regulation in MuSK-MG. Together, these findings support a role for epigenetic dysregulation of Th2-associated cytokines in the immunological environment associated with IgG4 subclass switch. To our knowledge, this is the first study investigating *IL4*, *IL10*, and *IL13* promoter DNA methylation in MuSK-MG.

## Introduction

Myasthenia gravis (MG) with antibodies against muscle-specific kinase (MuSK-MG) is a rare autoimmune disease of the neuromuscular junction characterized by fluctuating skeletal muscle weakness resulting from impaired synaptic transmission at the postsynaptic membrane. It is caused by antigen-specific autoantibodies that disrupt the interaction between MuSK and its binding partner low-density lipoprotein receptor-related protein 4 (Lrp4), leading to impaired clustering of acetylcholine receptors^1^. Notably, functional blockade is considered the primary pathogenic mechanism available to these antibodies, as they predominantly belong to the IgG4 subclass^1,2^.

IgG4 antibodies are typically observed after prolonged exposure, particularly in the context of allergic responses, and are a part of T helper 2 (Th2) immune response^3–5^. Their anti-inflammatory properties and association with improved allergic outcomes are attributed to several unique features, including reduced binding to Fcγ receptors and complement component C1q, dynamic Fab-arm exchange resulting in functional bispecificity, and high antigen affinity driven by extensive somatic hypermutation^6^. Despite these regulatory characteristics, IgG4 autoimmune disorders (IgG4-AID) have emerged as a distinct group of diseases defined by the presence of pathogenic, antigen-specific IgG4 autoantibodies^7^ currently comprising at least 29 recognized members^8^.

It remains unclear why the immune response in these conditions is skewed towards the otherwise immunologically inert IgG4 subclass. One hypothesis proposed by Koneczny et al^9^ suggests that individual predisposition to antigen-specific autoimmunity, driven by particular major histocompatibility complex (MHC) class II, antigen, and T-cell receptor interactions, may favor the development of an IgG4-dominant immune response. This concept is supported by strong genetic associations observed in MuSK-MG, including HLA-DQB1*05 and HLA-DRB1*14^10,11^.

Another possible explanation for the phenomenon is the involvement class-switching pathways, including an initial switch to IgG1/3 before undergoing a transition to IgG4^9^. Consistent with this, longitudinal observations in several IgG4-AID, such as autoimmune nodopathies, idiopathic thrombotic thrombocytopenic purpura and membranous nephropathy have demonstrated a shift in antigen-specific responses from IgG1 and/or IgG3 toward IgG4 in a subset of patients, supporting the existence of an intermediate IgG1/IgG3 predominant phase prior to IgG4 predominance^12–15^.

One potential mechanism underlying this subclass switching is the cytokine milieu, as increased production of Th2-associated cytokines, particularly interleukin (IL)-4, IL-10, and IL-13, plays a central role in IgG4 subclass switching^3–5^. Supporting the relevance of this pathway, increased expression of *IL4* and *IL10*, as well as elevated levels of these cytokines have been reported in IgG4-related disease, a fibroinflammatory condition characterized by IgG4-positive plasma cell infiltration and elevated total IgG4 levels^16,17^. Similarly, increased IL-10 levels have been detected in MuSK immunized mice^18^ and in the serum of patients with MuSK-MG^19^. In addition, elevated IL-10 levels have been reported in HLA-DRB1*14+ MuSK-MG patients^20^.

However, the mechanisms driving the dysregulated expression of these cytokines remain poorly understood. One potential explanation is epigenetic regulation of gene expression.

Epigenetic processes are defined as heritable changes in gene expression that occur without alterations in the underlying DNA sequence and include DNA methylation and histone modifications^21^. DNA methylation at CpG sites within promoter regions is typically associated with transcriptional repression. Transcriptional upregulation by hypomethylation in gene regulatory elements of interleukins has been shown in various disease contexts^22–26^, including *IL10* and *IL13* in systemic lupus erythematosus (SLE)^27^.

Therefore, we investigated promoter DNA methylation levels of *IL4*, *IL10*, and *IL13* in patients with MuSK-MG to determine whether epigenetic dysregulation of these Th2-associated cytokines may contribute to the IgG4-skewed immune response.

## Methods

### Cohort

The study cohort comprised patients with muscle-specific kinase antibody-positive myasthenia gravis (MuSK-MG), acetylcholine receptor antibody-positive myasthenia gravis (AChR-MG), and sex-matched healthy control individuals for comparative analyses. A total of 30 MuSK-MG patients were recruited at the MG Center of Pisa University Hospital, Pisa, Italy, and an additional 6 MuSK-MG patients were recruited at the Medical University of Vienna (MUW), Vienna, Austria, as well as the comparison groups including 12 healthy controls and 7 AChR-MG patients, as an IgG1 mediated autoimmune disease control.

Patient charts and reports were reviewed to collect data on the following variables: sex, age at sample acquisition, age at diagnosis, symptoms, immunosuppressive treatments and disease status at the time of sample collection. Disease activity at the time of sample collection was determined according to Wiendl et al^28^, for the purpose of statistical analyses, patients were categorized as having active disease or remission based on the documented clinical assessment.

### Ethics Approval

All experiments involving human participants or biological material were conducted in accordance with the Declaration of Helsinki and approved by the Ethics Commission of the Medical University of Vienna (EK Nr: 1442/2017). Additional approval for the study “Biological markers in patients with neurological or neuromuscular autoimmune diseases” (protocol HORIZON-MSCA-2022-DN-01//IgG4-TREAT) was obtained from the Ethics Committee of the Regione Toscana – Area Vasta Nord Ovest (04/09/2025). Written informed consent was obtained from all participants.

### Sample Preparation

Peripheral blood mononuclear cells (PBMCs) were isolated from study participants by density gradient centrifugation over Ficoll-Paque in LeucoSep tubes (Greiner Bio-One, Cat. No: 227288) according to the manufacturer’s instructions. Isolated PBMCs were washed with Dulbecco’s Phosphate Buffered Saline (DPBS), pooled, and cryopreserved at a concentration of 1 × 10^6^ cells/mL in a freezing medium consisting of 10% DMSO in fetal bovine serum (Sigma-Aldrich, Cat. No: F7524). Cryopreserved cells were stored at -80 °C until downstream analyses.

### Methylation-Sensitive High-Resolution Melting (MS-HRM) Analysis

Genomic DNA was isolated from PBMC samples using the QIAamp® DNA Blood Mini Kit (QIAGEN, Cat. No: 52304) according to the manufacturer’s instructions and quantified with a NanoDrop ND 2000c spectrophotometer (Thermo Fisher Scientific, Milan, Italy). For DNA methylation analysis, 200 ng of DNA from each sample was treated with sodium bisulfite using the EpiTect® Bisulfite Kit (QIAGEN, Milan, Italy), which converts unmethylated cytosines to uracil while leaving methylated cytosines unchanged.

The methylation status of *IL4*, *IL10* and *IL13* gene promoters was assessed using the Methylation-Sensitive High-Resolution Melting (MS-HRM) technique, following a protocol adapted from Stoccoro et al, 2023^29^. CpG-rich regions within the promoter regions of each gene were identified using Methprimer software^30^, and MS-HRM primers were designed accordingly (**Supplementary Table 1**). The experiments were performed using a CFX96 Real-Time PCR detection system (Bio-Rad, Milan, Italy). For each gene, mean methylation values were calculated across all interrogated CpG sites within the selected region. Standard DNA samples with defined methylation levels (0%, 12.5%, 25%, 50%, 75%, and 100%; Qiagen, Milan, Italy) were included in each assay to calibrate and quantify methylation in the test samples, following the protocol described previously ^31^. All measurements were performed in duplicate to ensure reliability and reproducibility.

### Quantitative Real-Time PCR

Total RNA was isolated from PBMC (1 × 10^6^ cells) using the RNeasy Total RNA Kit (Qiagen, Cat. No: 74104) and then reverse transcribed using High-Capacity RNA-to-cDNA™ Kit (Thermo Fisher, Cat No: 4387406). Briefly, RNA was quantified and either used directly or diluted to a standardized concentration prior to reverse transcription. Reactions were prepared in a total volume of 20 µl according to the manufacturer’s recommendations and subjected to reverse transcription at 25°C for 5 min, 46°C for 20 min, followed by enzyme inactivation at 95°C for 1 min and held at 0°C. Before qPCR, cDNA was diluted 1:10 in nuclease-free water.

Primer pairs for *IL4*, *IL10*, *IL13*, and *GAPDH* were designed using Primer-BLAST (NCBI) based on reference mRNA sequences obtained from GenBank. Primer specificity was assessed in silico during primer design. Primer sequences are listed in **Supplementary Table 2**.

Quantitative PCR was performed using SYBR Green chemistry in a total reaction volume of 10 µl containing 1.0 µl cDNA, 5.0 µl SsoAdvanced SYBR Supermix (Bio Rad, Cat No:1725272), 0.2 µl each forward and reverse primer, and 3.6 µl nuclease-free water. Amplification was carried out on an AriaMx Real-Time PCR System (Agilent Technologies) under the following cycling conditions: initial denaturation at 95°C for 30 s, followed by 40 cycles of denaturation at 95°C for 15 s and annealing/extension at 60°C for 30 s. Melting curve analysis (95°C for 15 s, 60°C for 30 s, and 95°C for 15 s) was performed after amplification to confirm product specificity. Relative expression of *IL4*, *IL10*, and *IL13* was normalized to *GAPDH* and calculated using the 2^×-ΔΔCt^ method^32^.

### IgG Subclass Analysis by Flow Cytometry

Antigen-specific IgG subclass levels were measured using a HEK293 cell–based assay adapted for flow cytometry from Koneczny et al, 2013^1^. Cells were transfected with a plasmid vector (pCMV6-AC-GFP) encoding full-length human MuSK (NM_005592, Origene, Cat. No: RG212253). The following day, 1×10⁵ cells per well were incubated with patient or control serum (1:40 dilution) in blocking medium (DMEM supplemented with 10% normal donkey serum, 1% bovine serum albumin and 5 mM HEPES) for 20 minutes at 4°C. After washing and fixation with 4% paraformaldehyde (Thermofisher Scientific, Cat. No: J61899), cells were stained with biotinylated anti-human IgG1 (8c/6-39, Sigma Aldrich, Cat. No: I2513), IgG2 (HP-6014, Sigma Aldrich, Cat. No: I5635), IgG3 (HP-6050, Sigma Aldrich, Cat. No: I7260), or IgG4 (HP-6025, Sigma Aldrich, Cat. No: I7385) antibodies (1:500). Cells were then washed with PBS and incubated with Cy3-conjugated Goat Anti-Mouse IgG (H+L) (Jackson Immuno Research, Cat. No: 115-165-166, 1:750). Cells were analyzed by flow cytometry (Cytoflex LX), with GFP expression distinguishing transfected from untransfected cells. Specific IgG subclass binding was calculated as the difference in median Y585-PE fluorescence intensity (ΔMFI) between transfected and untransfected cells. ΔMFI values were corrected for serum dilution and normalized within each experiment by dividing each sample by the mean ΔMFI of four positive control samples included in the same experiment. The resulting normalized ratios were then multiplied by the overall mean ΔMFI of all positive control measurements across experiments to reduce inter-assay variability.

### Total IgG4 Measurement by ELISA

Ninety-six–well flat-bottom plates were coated with 50 µL/well of anti-human IgG4 antibody (Invitrogen, cat. #A10651) diluted 1:1000 in phosphate-buffered saline (PBS) and incubated overnight at 4 °C. The plates were then washed six times with PBS containing 0.05% Tween-20 and blocked with 4% non-fat dry milk in PBS for 1 h at 37 °C. Following, samples and commercially available humanized IgG4 monoclonal antibody (natalizumab; Tysabri®) as standards diluted in PBS containing 1% bovine serum albumin (BSA) and 0.02% Tween-20 were added and incubated for 1 h at 37 °C. Plates were washed six times, and an F(ab’)₂ anti-human IgG Fcγ antibody conjugated to horseradish peroxidase (HRP), diluted 1:20,000 in 1% BSA and 0.02% Tween-20 in PBS, was added and incubated for 1 h at 37 °C. After a final washing step, substrate solution was added and incubated at room temperature for 20 min. The reaction was stopped with 2 M H₂SO₄, and absorbance was measured at 450 nm.

Serum samples were analyzed at 3 serial dilutions, each in duplicate and accepted when coefficient of variation was <15%.

### Statistical Analysis

Statistical analyses were performed using IBM SPSS Statistics version 29.0.2.0 (IBM Corp., Armonk, NY, USA) and GraphPad Prism version 10.6.1 (GraphPad Software, Boston, MA, USA). Normality was assessed using the Shapiro-Wilk test. As methylation data were not normally distributed, group differences were analyzed using the Kruskal-Wallis test followed by Dunn’s multiple comparisons test with Bonferroni adjustment. Exploratory multivariable linear regression models were performed to evaluate the independent effects of diagnostic group, age, sex, disease phase, and treatment on promoter DNA methylation levels.

Associations between methylation levels and serological parameters were assessed using Spearman’s rank correlation. Two-sided p-values ≤0.05 were considered statistically significant.

## Results

The study cohort comprised 36 patients with MuSK-MG (29 females, 7 males, median age 53 years), 7 patients with AChR-MG (4 females, 3 males, median age 50 years), and 12 healthy controls (9 females, 3 males, median age 33 years). At the time of sampling, 19 MuSK-MG patients were in remission and 17 had active disease. Regarding treatment status, 6 MuSK-MG patients were not receiving active immunosuppressive therapy. The remaining patients were treated with prednisolone (n = 29), with additional treatment including azathioprine (n = 1), methotrexate (n = 1), or rituximab (n = 3); one patient received methotrexate monotherapy (**Table 1**).

**Table 1:** Demographic and clinical characteristics of the study population.

| Characteristic | Healthy controls (n=12) | AChR-MG (n=7) | MuSK-MG (n=36) |
| --- | --- | --- | --- |
| Age, median (IQR), years | 33 (30-43) | 50 (45-80) | 53 (39-69) |
| Female, n (%) | 9 (75%) | 4 (57.1%) | 29 (80.6%) |
| Male, n (%) | 3 (25%) | 3 (42.9%) | 7 (19.4%) |
| Disease phase: remission, n (%) | — | 2 (28.5%) | 19 (52.7%) |
| Disease phase: active, n (%) | — | 5 (71.4%) | 17 (47.3%) |
| No active treatment, n (%) | — | 1 (14.3%) | 6 (16.7%) |
| Prednisolone, n (%) | — | 5 (71.4%) | 29 (80.5%) |
| Azathioprine, n (%) | — | 4 (57.1%) | 1 (2.8%) |
| Methotrexate, n (%) | — | 1 (14.3%) | 2 (5.6%) |
| Rituximab, n (%) | — | — | 3 (8.3%) |
| Max disease severity MGFA I, n (%) | — | — | 4 (11.1%) |
| Max disease severity MGFA II, n (%) | — | 2 (28.6%) | 18 (50%) |
| Max disease severity MGFA III, n (%) | — | 4 (57.1%) | 9 (25%) |
| Max disease severity MGFA IV, n (%) | — | 1 (14.3%) | 3 (8.3%) |
| Max disease severity MGFA V, n (%) | — | — | 2 (5.6%) |

Among AChR-MG patients, 2 patients were in remission, and 5 had active disease. One patient was not receiving active treatment, while 5 patients were treated with prednisolone, of whom 4 additionally received methotrexate. One patient received methotrexate monotherapy.

Significant differences in promoter methylation were observed for *IL4* and *IL13*, but not IL10 (**Table 2**, **Figure 1**). Pairwise comparisons demonstrated reduced *IL4* promoter methylation in MuSK-MG compared with both AChR-MG and healthy controls, while *IL13* promoter methylation was reduced only compared with healthy controls. No significant differences were detected for *IL10*.

**Figure 1.**
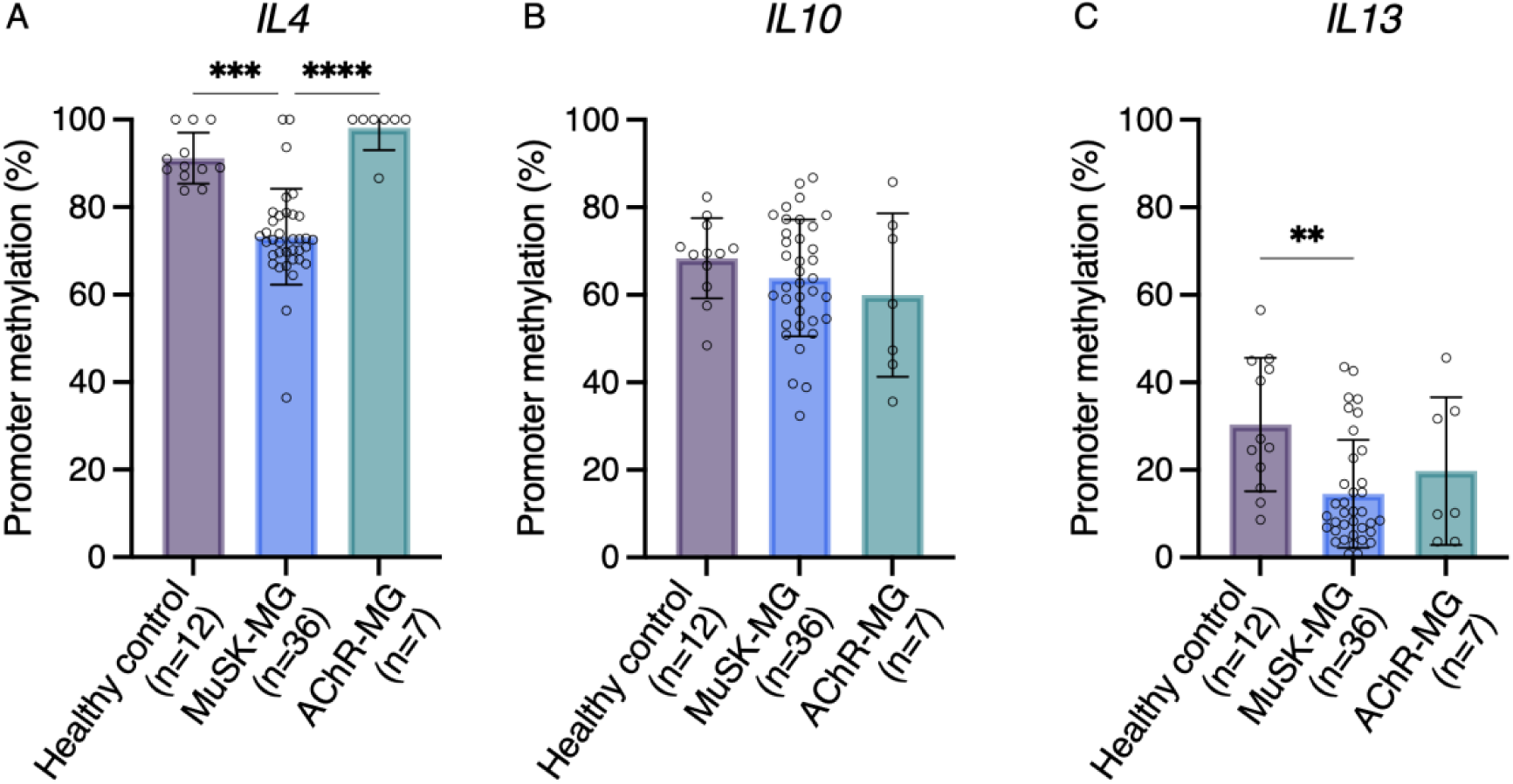
MuSK-MG patients exhibit reduced promoter DNA methylation of *IL4* and *IL13*, while *IL10* promoter methylation remains unchanged. Promoter methylation percentages were compared between healthy controls, AChR-MG patients, and MuSK-MG patients. Bars represent median methylation values and error bars indicate the interquartile range (25th–75th percentile). Individual data points are shown. Group differences were analyzed using the Kruskal–Wallis test followed by Bonferroni-adjusted pairwise comparisons. *IL4* and *IL13* promoter methylation levels were significantly reduced in MuSK-MG patients compared with healthy controls (adjusted p < 0.001and p = 0.004, respectively) and, for *IL4*, also compared with AChR-MG patients (adjusted p < 0.001). No significant differences were observed for *IL10* promoter methylation.

**Figure 2.**
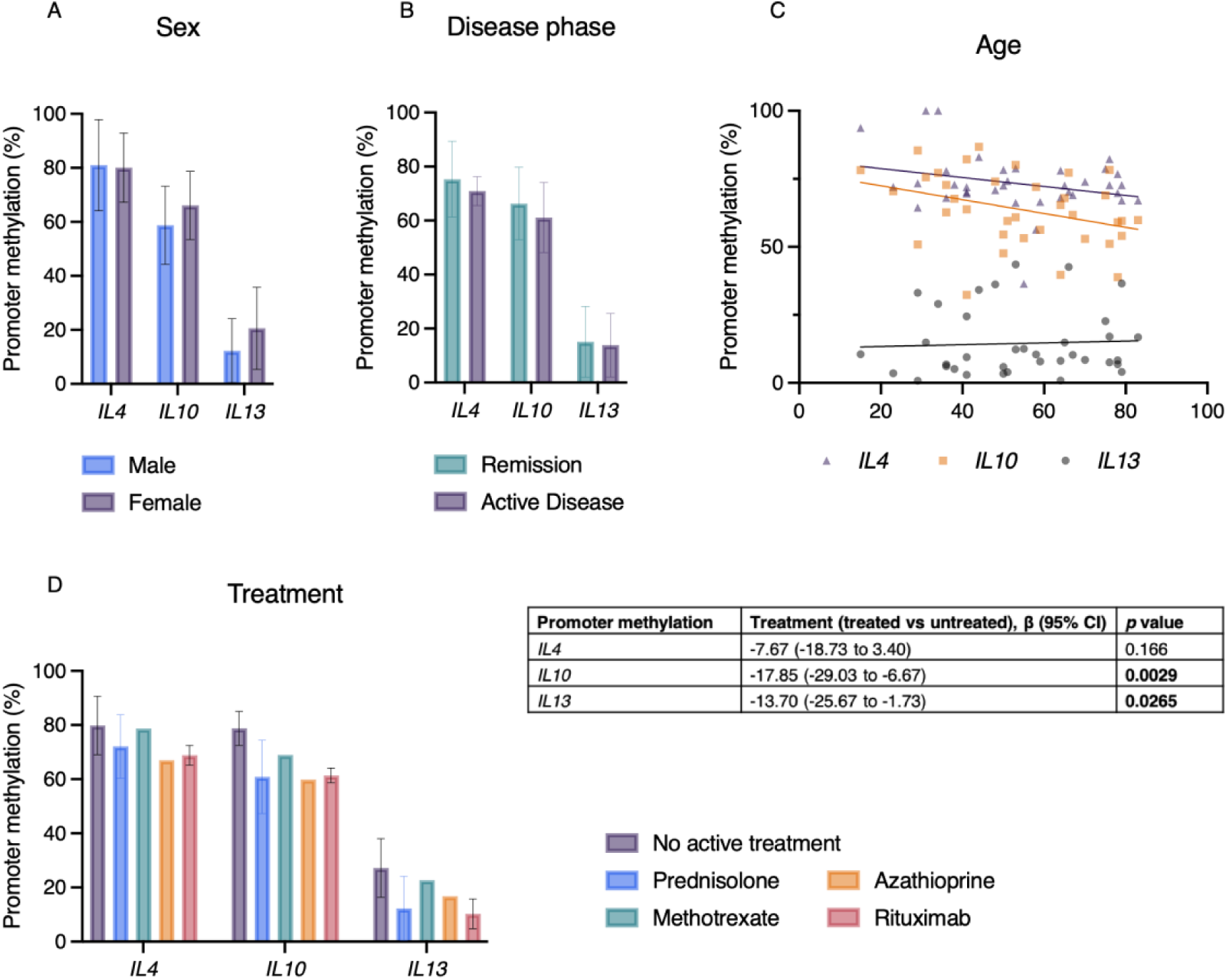
Treatment at sampling is independently associated with *IL10* and *IL13* promoter methylation in MuSK-MG patients. Promoter methylation levels were evaluated according to sex (A), disease phase (B), age (C), and treatment status (D). Bars represent mean values ± standard error of the mean. Age is presented as a scatter plot with a simple linear fit for visualization purposes. Independent associations between clinical variables and promoter methylation were assessed using multivariable linear regression analyses (Supplementary Table 3). Treatment at sampling was independently associated with lower *IL10* and *IL13* promoter methylation, whereas no independent associations were observed for *IL4*.

**Table 2:** Summary of group comparisons and correlation analyses of IL4, IL10, and IL13 promoter methylation.

| Parameter | Comparison | Statistical test | Effect size | 95% CI | Adjusted *p* value |
| --- | --- | --- | --- | --- | --- |
| **IL4 promoter methylation** | MuSK-MG vs AChR-MG | Kruskal–Wallis + Bonferroni | 72.56% (IQR 68–78) vs 100% (IQR 100–100) | — | <0.001 |
|  | MuSK-MG vs Healthy controls | Kruskal–Wallis + Bonferroni | 72.56% (IQR 68–78) vs 89.19% (IQR 88–98) | — | <0.001 |
|  | AChR-MG vs Healthy controls | Kruskal–Wallis + Bonferroni | 100% (IQR 100–100) vs 89.19% (IQR 88–98) | — | 1.000 |
| **IL10 promoter methylation** | MuSK-MG vs AChR-MG | Kruskal–Wallis + Bonferroni | 63.26% (IQR 54–75) vs 57.99% (IQR 44–76) | — | ≥0.756 |
| | MuSK-MG vs Healthy controls | Kruskal–Wallis + Bonferroni | 63.26% (IQR 54–75) vs 69.50% (IQR 63–75) | — | $\geq 0.756$ |
| | AChR-MG vs Healthy controls | Kruskal–Wallis + Bonferroni | 57.99% (IQR 44–76) vs 69.50% (IQR 63–75) | — | $\geq 0.756$ |
| ***IL13* promoter methylation** | MuSK-MG vs AChR-MG | Kruskal–Wallis + Bonferroni | 9.89% (IQR 6–21) vs 10.23% (IQR 4–33) | — | 1.000 |
|  | MuSK-MG vs Healthy controls | Kruskal–Wallis + Bonferroni | 9.89% (IQR 6–21) vs 26.11% (IQR 17–45) | — | **0.004** |
|  | AChR-MG vs Healthy controls | Kruskal–Wallis + Bonferroni | 10.23% (IQR 4–33) vs 26.11% (IQR 17–45) | — | 0.295 |
| ***IL4* promoter methylation** | *IL4* relative mRNA expression | Spearman rank correlation | $\rho = -0.7162$ | -0.8282 to -0.5492 | **$p = <0.0001$** |
| ***IL10* promoter methylation** | *IL10* relative mRNA expression | Spearman rank correlation | $\rho = -0.2046$ | -0.4543 to -0.07491 | $p = 0.1378$ |
| ***IL13* promoter methylation** | *IL13* relative mRNA expression | Spearman rank correlation | $\rho = -0.2639$ | -0.5027 to -0.01227 | $p = 0.0538$ |
| ***IL4* promoter methylation** | Total IgG4 | Spearman rank correlation | $\rho = -0.03296$ | -0.3664 to -0.3080 | $p = 0.8486$ |
| ***IL10* promoter methylation** | Total IgG4 | Spearman rank correlation | $\rho = -0.1734$ | -0.4827 to 0.1743 | $p = 0.3118$ |
| ***IL13* promoter methylation** | Total IgG4 | Spearman rank correlation | $\rho = -0.1429$ | -0.4583 to 0.2045 | $p = 0.4058$ |
| ***IL4***<br>**promoter**<br>**methylation** | Anti-MuSK<br>IgG4 | Spearman<br>rank<br>correlation | $\rho =$<br>0.09004 | -0.2552 to 0.4149 | $p =$<br>0.6015 |
| ***IL10***<br>**promoter**<br>**methylation** | Anti-MuSK<br>IgG4 | Spearman<br>rank<br>correlation | $\rho =$<br>-0.2175 | -0.5171 to 0.1295 | $p =$<br>0.2025 |
| ***IL13***<br>**promoter**<br>**methylation** | Anti-MuSK<br>IgG4 | Spearman<br>rank<br>correlation | $\rho =$<br>-0.2138 | -0.5142 to 0.1333 | $p =$<br>0.2106 |

To assess whether promoter DNA methylation was associated with clinical characteristics within the MuSK-MG cohort, multivariable linear regression analyses were performed including age, sex, treatment at sampling, disease phase, disease duration, and MGFA score (**Supplementary Table 3**). No independent associations between *IL4* promoter methylation and any of the investigated clinical variables were observed. Likewise, *IL10* and *IL13* promoter methylation were not associated with age, sex, disease phase, disease duration, or MGFA score. However, treatment at sampling was independently associated with lower *IL10* (β = −17.85, 95% CI −29.03 to −6.67, *p* = 0.0029) and *IL13* (β = −13.70, 95% CI −25.67 to −1.73, *p* = 0.0265) promoter methylation.

Next, we quantified relative mRNA expression of the cytokine genes *IL4*, *IL10*, *IL13* using qPCR. Analysis revealed significantly higher expression of *IL4* (p = 0.0005), *IL10* (p = 0.0462) and *IL13* (p = 0.0002) in MuSK-MG patients compared with AChR-MG patients. Compared with healthy controls, *IL4* expression was significantly increased in MuSK-MG patients (p < 0.0001). For *IL10* and *IL13* expression levels were also increased compared with healthy controls, although they did not reach statistical significance (p = 0.1874 and 0.0739, respectively) (**Figure 3**). Fold changes relative to healthy controls (2^-ΔΔCt^) are plotted in **Supplementary Figure 1**.

**Figure 3.**
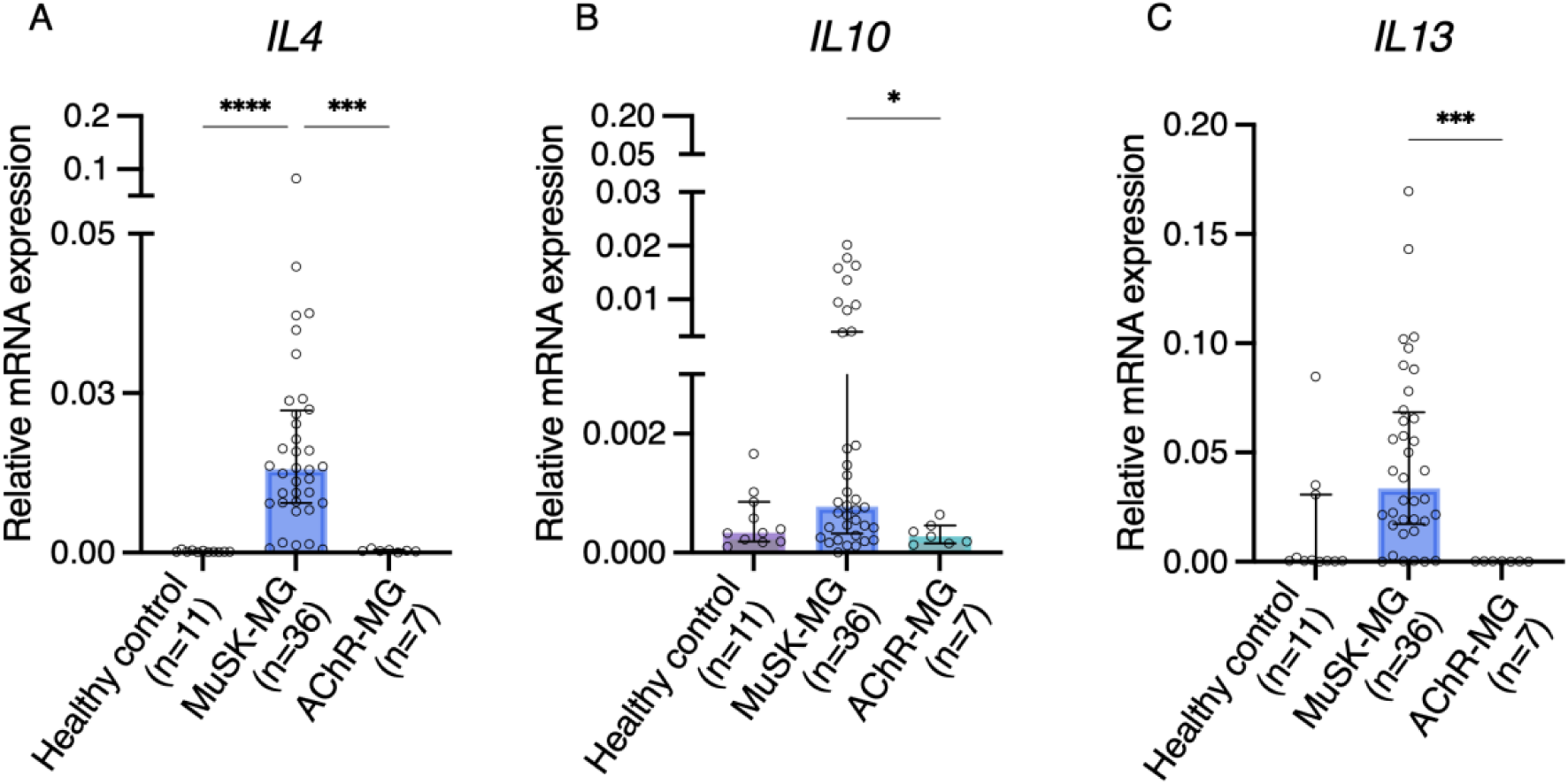
MuSK-MG patients show increased relative mRNA expression of *IL4, IL10* and *IL13*. Relative mRNA expression levels of *IL4*, *IL10*, and *IL13* were quantified by qPCR in MuSK-MG patients, AChR-MG patients, and healthy controls. MuSK-MG patients showed significantly increased expression of *IL4*, *IL10*, and *IL13* compared with AChR-MG patients. Compared with healthy controls, *IL4* expression was significantly increased in MuSK-MG patients, while *IL10* and *IL13* showed a similar pattern without reaching statistical significance. Data are presented as median with interquartile range (25th–75th percentile), with individual data points shown. Group differences were analyzed using the Kruskal– Wallis test followed by Bonferroni-adjusted pairwise comparisons. Significance is indicated as *p < 0.05, ***p < 0.001, ****p < 0.0001.

To evaluate the association between promoter methylation levels and relative mRNA expression levels, correlation analyses were performed for *IL4*, *IL10* and *IL13*. A significant inverse correlation was observed for *IL4* and for *IL13,* an inverse correlation was also noted, however it did not reach statistical significance (p = 0.0538). No correlation between *IL10* promoter methylation levels and mRNA expression was seen (**Table 2**, **Figure 4**).

**Figure 4.**
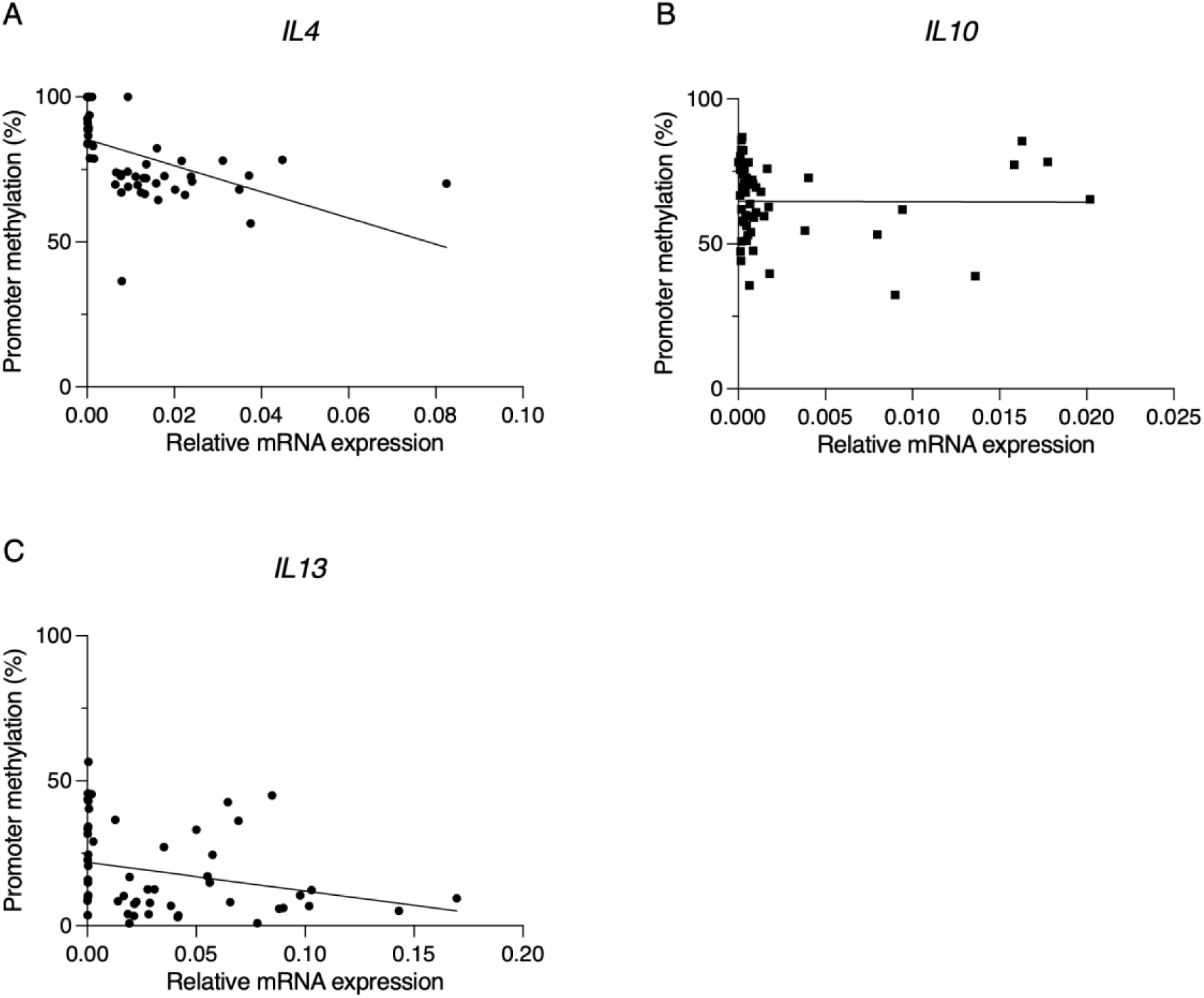
Promoter methylation levels of *IL4* and *IL13* are inversely correlated with mRNA expression levels. Scatter plots show the association between promoter DNA methylation levels and relative mRNA expression of *IL4*, *IL10*, and *IL13*. Each point represents an individual sample. Lines represent simple linear fits for visualization. An inverse correlation between promoter methylation and relative mRNA expression was observed for *IL4*, with a similar but non-significant trend for *IL13*, whereas no association was detected for *IL10*.

Finally, we assessed whether promoter DNA methylation levels were associated with serological characteristics. Spearman correlation analysis revealed no significant correlations between *IL4*, *IL10*, or *IL13* promoter methylation and total IgG4 concentrations or the percentage of IgG4 within anti-MuSK antibodies (**Table 2**, **Figure 5**). Anti-MuSK IgG4 antibodies also did not correlate with the relative mRNA expression of the selected cytokines (**Supplementary Figure 2**). Similarly, no correlations were observed between anti-MuSK IgG4 ΔMFI values and promoter methylation or relative mRNA expression levels of *IL4*, *IL10*, or *IL13* (**Supplementary Figure 3**). Additional analyses of the non-IgG4 subclasses likewise showed no significant correlations promoter methylation levels and the proportions of IgG1, IgG2, or IgG3 (**Supplementary Figure 4**).

**Figure 5.**
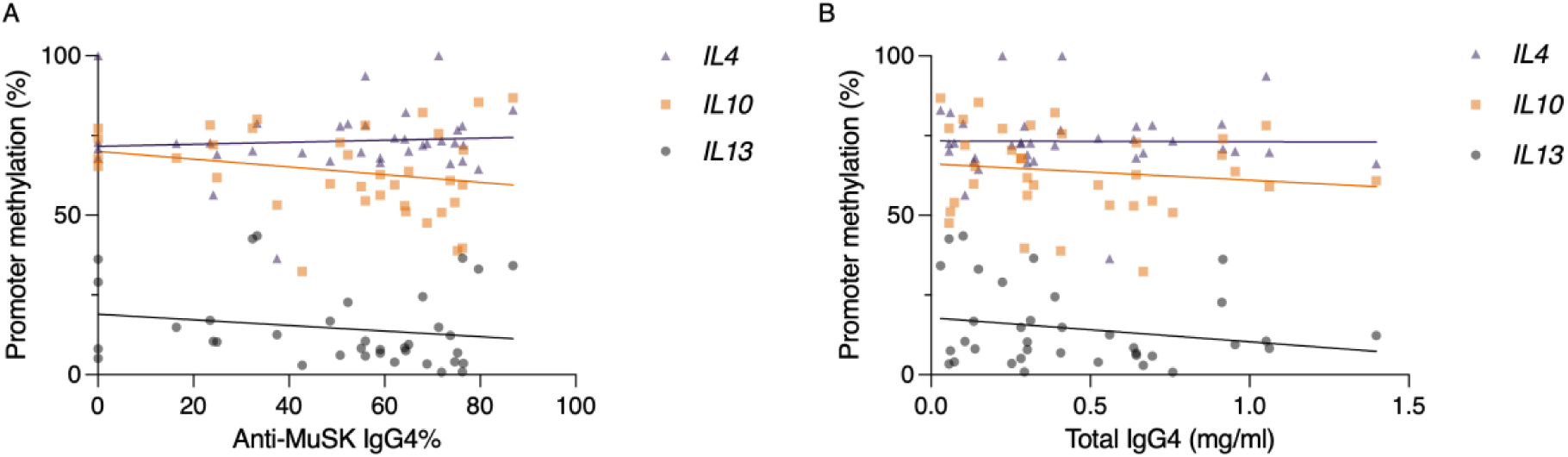
No association observed between anti-MuSK IgG4 percentages, total IgG4 concentrations and promoter DNA methylation levels. Scatter plots show *IL4*, *IL10*, and *IL13* promoter DNA methylation levels in relation to total IgG4 concentrations and the proportion of IgG4 within MuSK antibodies. Lines represent simple linear fits for visualization. Spearman correlation analysis revealed no significant associations between methylation levels and IgG4-related measures.

## Discussion

MuSK myasthenia gravis is a well-characterized IgG4-autoimmune disease; however, the mechanisms driving the predominance of this antibody subclass remain incompletely understood. Given the established role of IL-4, IL-10, and IL-13 in IgG4 class switching and emerging evidence that cytokine pathways are subject to epigenetic regulation, we investigated whether promoter methylation and gene expression levels of these cytokines are altered and associated with IgG4 antibody levels in this disease.

In the present study, we identified significantly lower promoter methylation levels of *IL13* and *IL4* in MuSK-MG patients compared with healthy controls, while *IL4* methylation was additionally reduced compared with AChR-MG disease controls. In contrast, no significant differences were observed for *IL10* promoter methylation. We also observed increased expression of *IL4*, *IL10*, and *IL13* in MuSK-MG patients compared with AChR-MG patients. *IL4* expression was significantly upregulated in MuSK-MG patients compared with healthy controls and showed a significant inverse correlation with promoter methylation levels, supporting a potential relationship between epigenetic regulation and transcriptional activity. A trend for inverse association was also observed for *IL13*, but did not reach statistical significance.

These findings are in line with previous reports of elevated *IL4* and *IL10* expression and/or cytokine levels in IgG4-RLD and MuSK-MG^16,17,19^ and hypomethylation of *IL13* regulatory regions in CD4+ T cells from SLE patients negatively correlating with *IL13* expression, supporting a functional role of methylation in regulating Th2-associated cytokine responses^33^.

The observed changes in IL-4 and IL-13 are highly relevant in the context of IgG4 biology, as they are key mediators of IgG4 and IgE class switching, and IL-10 has been proposed to tip the balance towards IgG4^4,34^. This epigenetically regulated cytokine milieu may reflect a tolerogenic immune state that normally develops during prolonged antigen exposure and is permissive for IgG4 predominance in MuSK-MG.

From a translational perspective, the stronger association observed for *IL4* is notable given the emerging interest in targeting IL-4 signaling pathways. The observation that *IL4* promoter hypomethylation was not associated with demographic or clinical variables further supports IL-4 dysregulation as a fundamental feature of MuSK-MG rather than a secondary consequence of disease characteristics or treatment. Blockade of IL-4 and IL-13 signaling has been shown to modulate IgG4 responses in other immune contexts, raising the possibility that cytokine-directed approaches may also influence subclass switching in IgG4-mediated autoimmunity^35,36^. Whether epigenetic regulation of these pathways could provide additional therapeutic opportunities remains to be investigated.

Interestingly, despite observing increased *IL10* mRNA expression in MuSK-MG patients, no differences in promoter methylation were detected. This finding differs from a previous study that showed hypomethylation of the same promoter region in rheumatoid arthritis^37^, this discrepancy may instead reflect the complex and context-dependent regulation of *IL10* expression, rather than absence of epigenetic regulation in MuSK-MG. Previous studies have demonstrated that *IL10* expression can be associated with DNA demethylation in specific regulatory regions outside the promoter, including STAT1- and STAT3-mediated transcriptional activation in Th1 cells^38^. Similarly, studies in IL-10-producing B cells demonstrated that acquisition of *IL10* expression was associated with differential methylation of conserved non-coding regulatory regions within the *IL10* locus rather than promoter methylation^39^. Therefore, the lack of difference in *IL10* promoter methylation in bulk PBMCs may indicate that disease-relevant epigenetic changes occur at alternative regulatory regions not captured in the present study.

In contrast to sex, disease phase, disease duration, maximum disease severity, and age, treatment status was independently associated with lower *IL10* and *IL13* promoter methylation. To our knowledge, an association between immunosuppressive treatment and promoter methylation of these cytokines has not previously been reported in MuSK-MG or other IgG4-AID. Previous studies have shown that corticosteroid exposure increases *IL10* mRNA expression in CD4+ T cells and is accompanied by widespread DNA demethylation^40^, supporting the concept that glucocorticoids exert epigenetic effects on immune cells. Likewise, corticosteroid-induced hypomethylation of the IL12B locus has been linked to reduced IL12B expression^41^, consistent with the established role of IL-12 in promoting Th1 responses, whereas reduced IL-12 signaling favors Th2 differentiation^42^.

Together, these findings suggest that corticosteroids may contribute to epigenetic remodeling of cytokine regulatory loci, potentially promoting the Th2-skewed immune environment characteristic of MuSK-MG.

This observation may be particularly relevant in MuSK-MG, where corticosteroids remain a cornerstone of therapy but patients frequently require higher doses and prolonged treatment compared with AChR-MG, particularly during tapering^43^. Although the cross-sectional design of the present study precludes causal inference, we cannot exclude that the reduced IL13 promoter methylation observed in MuSK-MG was influenced, at least in part, by immunosuppressive treatment rather than the disease itself. This interpretation is further limited by the lack of cumulative treatment duration and dose information.

We additionally did not observe significant correlations between promoter methylation levels and either total IgG4 concentrations or the proportion of IgG4 within MuSK-specific antibodies. Similarly, relative mRNA expression levels of *IL4*, *IL10*, and *IL13* were not associated with these serological measures. Although this may initially appear unexpected given the role of IL-4 and IL-13 in IgG4 class switching, circulating antibody levels represent the cumulative outcome of multiple immunological processes, as well as clinical factors. These include germinal center maturation, plasma cell differentiation, disease activity and treatment exposure especially B cell depleting therapies, such as rituximab^44–49^. Consequently, serum antibody levels may not directly reflect upstream epigenetic states or transcriptional profiles that contributed earlier to disease development.

Similarly, the absence of association with total IgG4 levels may not be surprising in the context of IgG4-AID. Previous work from our group demonstrated that total serum IgG4 concentrations are not consistently elevated across IgG4-AID, despite the predominance of pathogenic IgG4 autoantibodies^50^. This suggests that the epigenetic alterations identified here are unlikely to induce generalized IgG4 overproduction, but may instead contribute to antigen-specific IgG4 subclass switch.

Several limitations should be considered when interpreting these findings. Most importantly, methylation analyses were performed on bulk PBMC populations rather than isolated immune cell subsets. As PBMCs represent a highly heterogeneous mixture of lymphoid and myeloid cells, disease-relevant epigenetic changes occurring in specific cellular populations may have been diluted or masked. Although PBMC composition was not assessed in the present cohort, the concordance between reduced *IL4* and *IL13* promoter methylation and increased *IL4* and *IL13* transcription suggests that these findings are unlikely to be explained solely by differences in cellular composition. In addition, MS-HRM measures average methylation across the amplified region and does not provide CpG site-specific resolution.

The cross-sectional study design also precludes determination of whether the observed methylation changes represent a cause or a consequence of disease. Furthermore, complementary cytokine measurements were not available for the present cohort, precluding direct evaluation of the relationship between promoter methylation and circulating cytokine levels. The relatively small AChR-MG cohort may also have limited statistical power for disease-control comparisons. Finally, samples were obtained from two recruitment sites, and potential site-related differences in sample handling, storage, or DNA processing cannot be excluded. Future studies employing cell type- and CpG site-specific epigenetic approaches, particularly focusing on IgG4-producing B cells and plasmablasts, together with functional immune profiling, may provide greater mechanistic insight into the biological significance of the observed methylation patterns.

Collectively, the present study points toward a potential contribution of epigenetic dysregulation of *IL4* and *IL13* to the characteristic IgG4-dominant immune response in MuSK-MG. To our knowledge, this is the first study to characterize promoter DNA methylation of Th2-associated cytokines in MuSK-MG and to integrate these findings with cytokine gene expression, providing new insight into the epigenetic mechanisms that may be involved in IgG4-mediated autoimmunity. These results support further investigation of epigenetic regulation in IgG4-AID and highlight the need for cell type-specific studies integrating methylation, transcriptional, and functional immune analyses.

## Acknowledgements

The authors gratefully acknowledge the Core Facility Flow Cytometry of the Medical University of Vienna, particularly Prof. Andreas Spittler, for technical support and expertise. The authors also thank Dr. Verena Endmayr and Dr. Carmen Haider for their assistance. We sincerely thank all patients and healthy volunteers who participated in this study making this research possible.

## Data availability

Anonymized study data are available upon reasonable request from the corresponding author.

## Study Funding

This project is conducted within the IgG4-TREAT consortium, a Marie Skłodowska-Curie Actions doctoral network funded by the European Union (Grant Agreement No. 101119457).

## Disclosure

**CE** and **FF** were supported by the HORIZON Europe Marie Skłodowska-Curie Actions Doctoral Network *IgG4-TREAT* (Grant Agreement No. 101119457). **IK** was supported by the same program, as well as by a Hertha Firnberg project grant from the Austrian Science Fund (FWF; T996-B30) and the argenx project grant *Mya-DACH*. **MW** was supported by the argenx project grant *Mya-DACH*. **OK** received support to attend scientific conferences or meetings from Amgen, Biogen, Lilly, Novartis, Roche, Sanofi and UCB, speaker’s honoraria from Biogen and Roche, and project funding from ArgenX. The authors report no conflict of interest.

## Author Contributions

Conceptualization: CE, IK. Methodology: CE, AS, JH, FF, FC, IK. Investigation: CE, AS, ML, FS. Formal analysis: CE. Resources: VI, AC, MW, OK, RW, AS, TK, MG, FZ, RH, MM, IK. Data curation: CE. Writing (original draft): CE. Review & editing: CE, AS, FC, IK. Supervision: AS, FC, IK. Funding acquisition: IK.

## Notes

### Competing Interest Statement

The authors have declared no competing interest.

