## Supplementary data for "Epigenetic dysregulation of Th2 cytokine genes in MuSK myasthenia gravis and its modulation by immunosuppressive therapy"

### Supplementary Tables and Figures

**Supplementary Table 1.** The main characteristics of the designed primers for MS-HRM, including primer sequences, annealing temperature (Ta), amplicon size, number of CpG sites included in the analyzed amplicons, and the genomic location of the analyzed regions are reported.

| Gene | Primer Sequence | T_a_ | Amplicon Size | CpG sites | Accession Number and  Nucleotide Position |
| --- | --- | --- | --- | --- | --- |
| IL4 | F-GTTGTGTGTTTTGGATTGTTATTAATTAT | 61°C | 263 | 4 | NG_023252.1:  10363–10626 (reverse strand) |
|  | R-ATAAATCTCACCTCCCAACTACTTC |  |  |  |  |
| IL10 | F-TGGGGAAATTAAGGTTTAGAGATTTA | 56°C | 215 | 4 | NG_012088.1: 4518-4733 |
|  | R-TTTCCTAAAAAAAACAACTATTCTATAC |  |  |  |  |
| IL13 | F-GAGGAAGGGAGGTTTTAATTTTAG | 57.8°C | 290 | 20 | NC_000005.10: 132655840-132656130 |
|  | R-ACCCTAAAAAATAATCTCCAAAAAC |  |  |  |  |

**Supplementary Table 2**. The main characteristics of the designed primers for qPCR, including target genes, primer sequences, and corresponding GenBank reference accession numbers used for primer design, are reported.

| Gene | Primer Sequence (5'→3') | GenBank Accession Number |
| --- | --- | --- |
| IL4 | F-AACGGCTCGACAGGAACCT | NM_000589.4 |
|  | R-ACTCTGGTTGGCTTCCTTCCA |  |
| IL10 | F-CGGCGCTGTCATCGATTT | NM_000572.3 |
|  | R-TTAAAGGCATTCTTCACCTGCTC |  |
| IL13 | F-GGAGCTGGTCAACATCACCC | NM_002188.3 |
|  | R-CGTTGATCAGGGATTCCAGG |  |
| GAPDH | F-ATATTGTTGCCATCAATGACCC | NM_002046.7 |
|  | R-ATGACAAGCTTCCCGTTCTC |  |

**Supplementary Table 3.** Multivariable linear regression analysis of clinical variables associated with promoter DNA methylation in the MuSK-MG cohort. Regression models included age, sex, treatment at sampling (treated vs untreated), disease phase, disease duration, and MGFA score. Regression coefficients (β), 95% confidence intervals (CI), and corresponding p values are shown for *IL4*, *IL10*, and *IL13* promoter methylation.

| **Predictor** | ***IL4*** β  **95% CI** | **p value** | ***IL10*** β  **95% CI** | **p value** | ***IL13*** β  **95% CI** | **p value** |
| --- | --- | --- | --- | --- | --- | --- |
| Age | −0.118 (−0.389 to 0.153) | 0.378 | −0.166 (−0.418 to 0.087) | 0.190 | 0.034 (−0.236 to 0.305) | 0.797 |
| Sex (Female vs Male) | 0.143  (-10.75 to 11.04) | 0.979 | 2.742  (-8.264 to 13.75) | 0.613 | 2.684  (-9.100 to 14.47) | 0.644 |
| Treatment (treated vs untreated) | -7.666  (-18.73 to 3.401) | 0.166 | -17.85  (-29.03 to -6.67) | **0.0029** | -13.70  (-25.67 to -1.73) | **0.0265** |
| Disease phase (Active vs Remission) | -5.294  (-14.85 to 4.263) | 0.265 | -3.719  (-13.38 to 5.936) | 0.436 | -3.225  (-13.56 to 7.113) | 0.527 |
| Disease duration | -0.052  (-0.598 to 0.493) | 0.846 | -0.343  (-0.894 to 0.208) | 0.212 | 0.034  (-0.556 to 0.624) | 0.906 |
| MGFA II vs I | 9.440  (-5.328 to 24.21) | 0.200 | 2.063  (-12.86 to 16.98) | 0.779 | 7.544  (-8.429 to 23.52) | 0.341 |
| MGFA III vs I | 8.251  (-7.194 to 23.70) | 0.282 | 3.965  (-11.64 to 19.57) | 0.606 | 6.026  (-10.68 to 22.73) | 0.465 |
| MGFA IV vs I | 3.928  (-17.45 to 25.31) | 0.709 | -6.776  (-28.37 to 14.82) | 0.525 | 0.810  (-22.31 to 23.93) | 0.943 |
| MGFA V vs I | 3.997  (-16.88 to 24.87) | 0.697 | -0.388  (-21.48 to 20.70) | 0.970 | -1.582  (-24.16 to 21.00) | 0.887 |

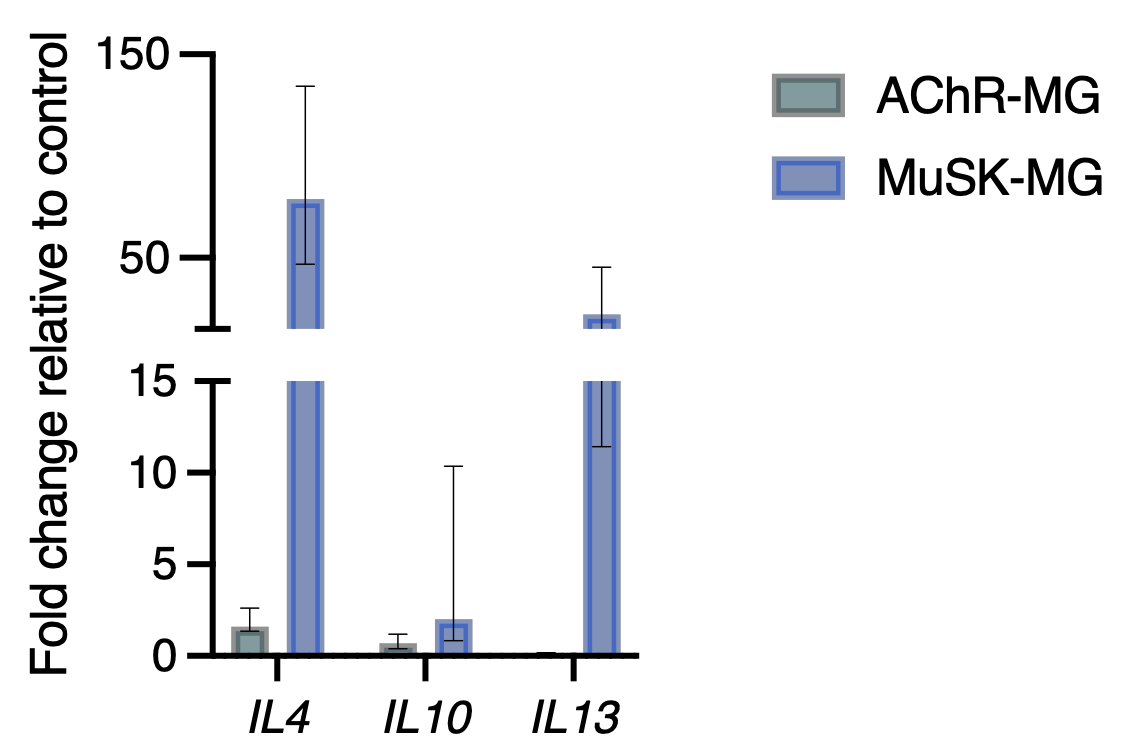

**Supplementary Figure S1.** Relative cytokine gene expression in MuSK-MG and AChR-MG presented as fold change relative to healthy controls.

Relative mRNA expression of IL4, IL10, and IL13 was quantified by qPCR and calculated using the 2^-ΔΔCt^ method. Expression levels were normalized to the housekeeping gene GAPDH and expressed as fold change relative to healthy controls used as the calibrator group. Data are presented as median with interquartile range (25th–75th percentile), with individual data points shown.

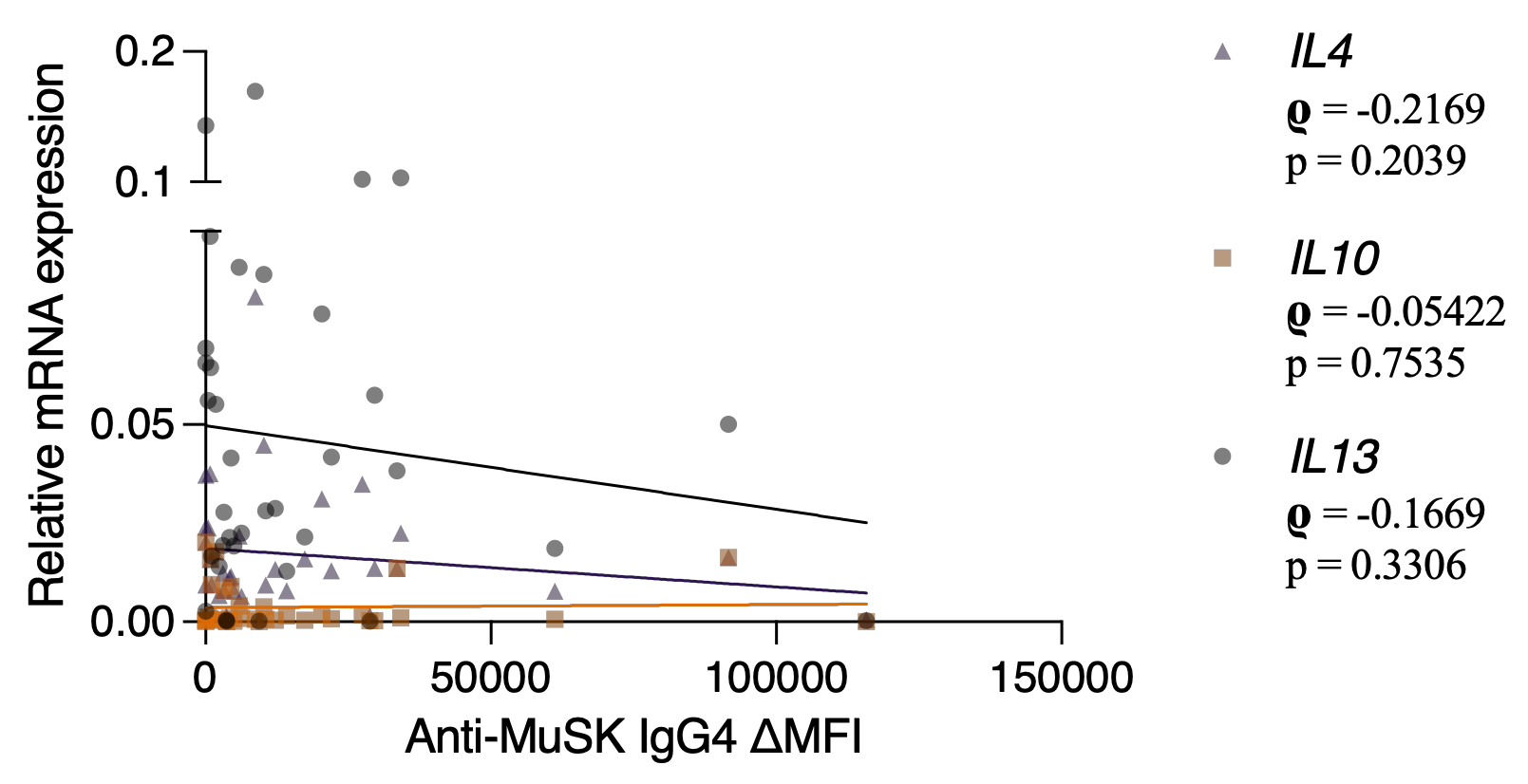

**Supplementary Figure S2. Association between relative mRNA expression and anti-MuSK IgG4 antibody percentages.**

Scatter plots illustrate the relationship between **relative mRNA expression** for *IL4*, *IL10*, *IL13* and percentage of IgG4 within anti-MuSK specific antibodies. Lines represent simple linear fits for visualization. Spearman correlation analysis revealed no significant associations (p>0.05).

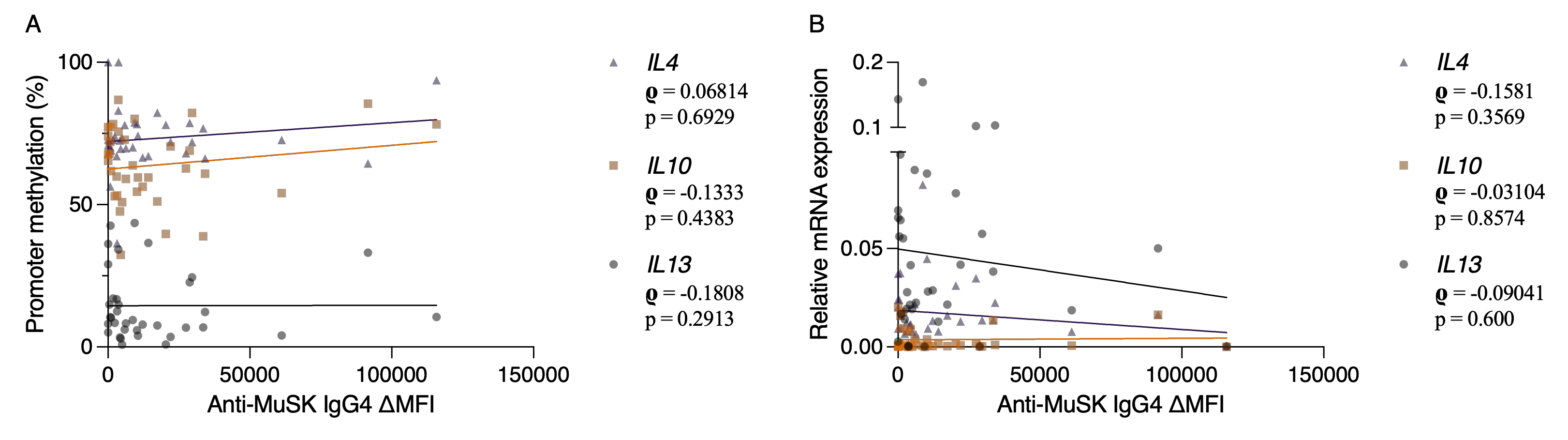

**Supplementary Figure S3. Association between promoter methylation, relative mRNA expression levels and anti-MuSK IgG4 antibody levels.**

Scatter plots illustrate the relationship between promoter methylation (A) or **relative mRNA expression (B)** for *IL4*, *IL10*, *IL13* and levels of IgG4 anti-MuSK specific antibodies. Lines represent simple linear fits for visualization. Spearman correlation analysis revealed no significant association (p>0.05).

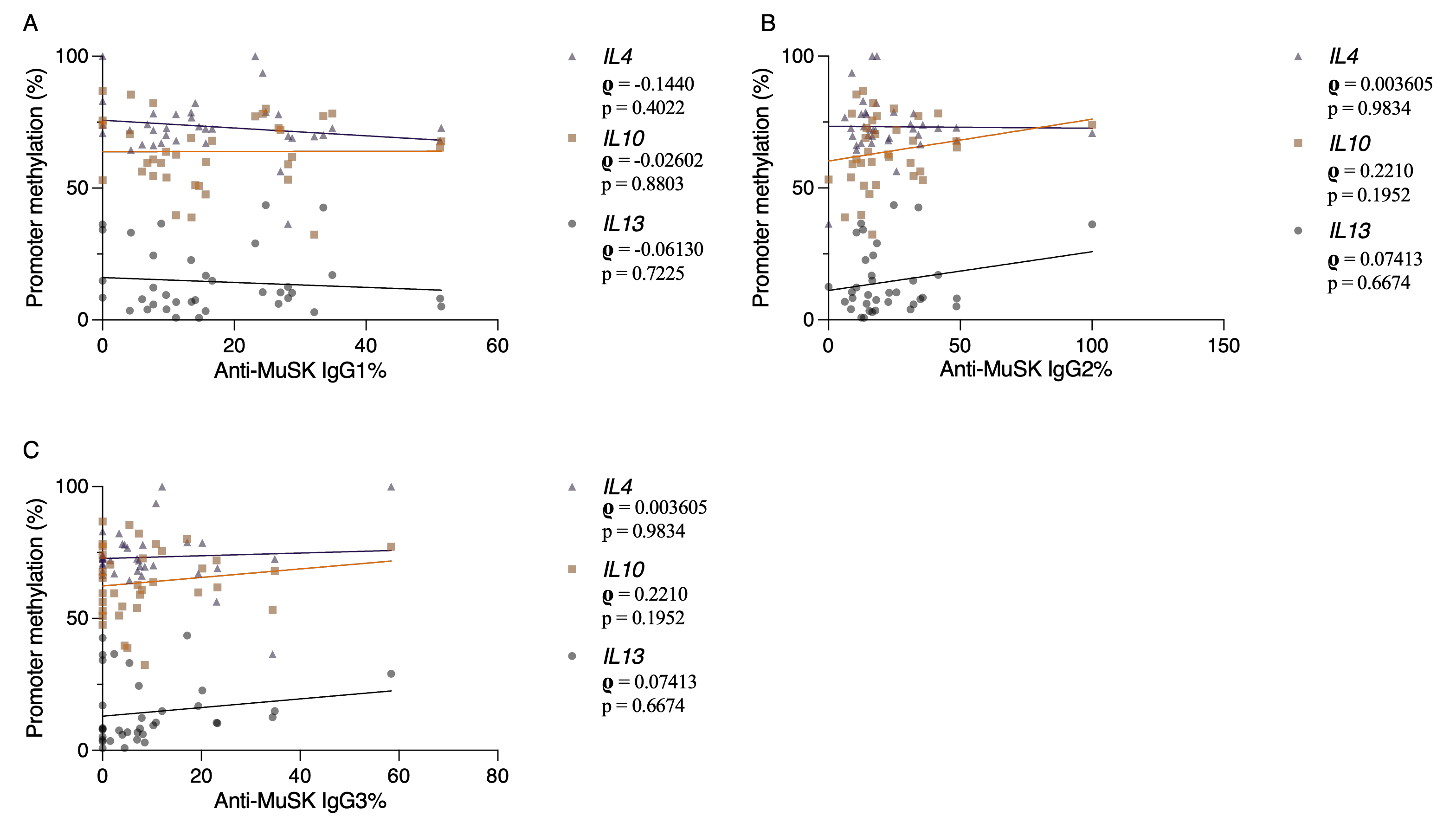

**Supplementary Figure S4. Association between promoter methylation levels and anti-MuSK IgG1,2,3 antibody percentages.**

Scatter plots showing the relationship between promoter DNA methylation of ***IL4***, ***IL10***, and ***IL13*** and the corresponding serum proportions of **IgG1 (A), IgG2 (B),** and **IgG3 (C).** Correlation coefficients (Spearman's ρ) and two-tailed p values are indicated on each graph. Linear fits are shown for visualization only.
